# Inferring chromosomal instability from copy number aberrations as a measure of chromosomal instability across human cancers

**DOI:** 10.1101/2023.05.24.542174

**Authors:** Sasha Taluri, Vishal H. Oza, Tabea M. Soelter, Jennifer L. Fisher, Brittany N. Lasseigne

## Abstract

**Background:** Cancer is a complex disease that is the second leading cause of death in the United States. Despite research efforts, the ability to manage cancer and select optimal therapeutic responses for each patient remains elusive. Chromosomal instability (CIN) is primarily a product of segregation errors wherein one or many chromosomes, in part or whole, vary in number. CIN is an enabling characteristic of cancer, contributes to tumor-cell heterogeneity, and plays a crucial role in the multistep tumorigenesis process, especially in tumor growth and initiation and in response to treatment.

**Aims:** Multiple studies have reported different metrics for analyzing copy number aberrations as surrogates of CIN from DNA copy number variation data. However, these metrics differ in how they are calculated with respect to the type of variation, the magnitude of change, and the inclusion of breakpoints. Here we compared metrics capturing CIN as either numerical aberrations, structural aberrations, or a combination of the two across 33 cancer data sets from The Cancer Genome Atlas (TCGA).

**Methods and results:** Using CIN inferred by methods in the CINmetrics R package, we evaluated how six copy number CIN surrogates compared across TCGA cohorts by assessing each across tumor types, as well as how they associate with tumor stage, metastasis, and nodal involvement, and with respect to patient sex.

**Conclusions:** We found that the tumor type impacts how well any two given CIN metrics correlate. While we also identified overlap between metrics regarding their association with clinical characteristics and patient sex, there was not complete agreement between metrics. We identified several cases where only one CIN metric was significantly associated with a clinical characteristic or patient sex for a given tumor type. Therefore, caution should be used when describing CIN based on a given metric or comparing it to other studies.

## 1. Introduction

Genomic instabilities, molecular signatures of gross genomic alterations, are enabling characteristics of cancer etiology and pathogenesis (1,2). They result in chromosomal breakages and rearrangements that can develop into chromosomal instability (CIN). CIN is primarily caused by defective cell cycle quality control mechanisms, including an elevated rate of segregation errors altering chromosomal content (3), and manifests as either numerical aberrations, structural aberrations, or a combination of the two. Numerical aberrations are whole chromosomal aberrations that lead to the loss of heterozygosity (i.e., where a chromosomal region is lost in one copy for a diploid genome) and variability in gene dosage effects. This can result in a phenotype that is a consequence of chromosome-wide altered expression patterns. Structural aberrations are sub-chromosomal and can lead to the fusion of gene products or amplified and/or deleted genes, specifically impacting the genes of the affected chromosomal regions (3). At the molecular level, CIN has been shown to represent distinct etiologies, promote disease progression, and metastases. It has also been associated with patient prognosis, drug efficacy, and drug resistance across many cancers (4–7). As CIN has been associated with poor patient outcomes in some cancers but improved survival in others (8,9), CIN appears to have cell- and tissue-specific consequences associated with the originating tissue and tumor site (10,11). For example, while CIN has been shown to have a non-monotonic relationship with patient outcome in ER–/ERBB2– breast, gastric, ovarian, squamous non-small cell lung carcinomas (i.e., patients with the lowest or highest quartile of CIN have a significantly improved hazard ratio) (12), it has been associated with poorer prognosis in diffuse large B-cell lymphoma patients (13). Therefore, CIN is a promising biomarker of patient prognosis and drug response but requires tissue-specific evaluation.

Several CIN scores have been proposed for analyzing copy number aberrations (i.e., deletions or amplifications of segments of the genome) as surrogates of chromosomal instability. Still, these scores represent various aspects of CIN and have been associated with different clinical and biological phenotypes across several cancer types (14–17). Therefore, assessing copy number aberration CIN surrogates across cancer types and tissue backgrounds is critical to evaluate the role of CIN in cancer etiology and progression. In addition, it enables the comparison of key results across studies and the identification of robust scores for future biomarker development. We recently published an R package, CINmetrics (18), for calculating six different copy number aberration CIN surrogate metrics, and here apply it to The Cancer Genome Atlas (TCGA). These metrics include total aberration index (TAI) (14), modified TAI, copy number abnormality (CNA) (15), number of break points (16), altered base segments (17), and fraction genome altered (FGA).

Specifically, they differ based on their ability to detect structural, numerical, or whole genome instability (discussed in depth in (18)). In this study, we aim to provide a comprehensive comparison of these metrics across a wide range of cancer types and with respect to clinical characteristics (tumor stage, node, and metastasis) and patient sex. We determine these CIN metric scores across 33 cancers and 22,629 samples from 11,124 patients and provide comparison statistics to evaluate how CIN metric scores vary across and within different cancers based on CIN classification (numerical, structural, or global), clinical characteristics, and patient sex.

## 2. Methods and Statistical Analysis

We downloaded masked copy number variation (CNV) (Affymetrix SNP 6.0 array) data and associated patient clinical data from all 33 projects in the TCGA portal using the ‘TCGAbiolinks’ R package (2.18.0) (19) in August 2021 using R (Version 4.0.4) and RStudio (Version 1.4.1106) locally and stored within a CSV file. We also downloaded TCGA Level 3, normalized and aggregated RNA-seq count data in November 2021 from all 33 projects in the TCGA portal using the ‘TCGAbiolinks’ R package (Version 2.22.1) using R (Version 4.0.2) and RStudio (Version 1.1.463) with University of Alabama at Birmingham’s High-Performance Computing Cluster, Cheaha. All analyses associated with this paper are on GitHub (https://github.com/lasseignelab/CINmetrics_Cancer_Analysis) and available at https://zenodo.org/record/7942543#.ZGPu8OzMJ4A.

With the CINmetrics R package (Version 0.1.0), we calculated each CIN metric for the 22,629 non-tumor and tumor samples (18). CINmetrics analyzes six different copy number aberration CIN surrogate metrics from masked CNV data as previously described (14–17). The mathematical formulas for those metrics are described in detail in Oza, et al. 2023, but briefly, those metrics are TAI (Total Aberration Index), Modified TAI, CNA (Copy Number Abnormality), Base Segments (i.e., the number of altered bases), Break Points (i.e., the number of break points), and FGA (Fraction of Genome Altered). Each metric defines chromosomal instability (CIN) by calculating numerical and/or structural aberrations as described in Oza et. al. (18) and can be grouped as numerical scores (Base Segments, FGA), structural scores (Break Points and CNA), and overall scores (TAI and Modified TAI). CNA and Break Points both consider the segmental abnormalities of the chromosome, but CNA requires that adjacent segments have a difference in segmentation mean values. TAI and Modified TAI can both be interpreted as the absolute deviation from the normal copy number state averaged over all genomic locations, where Modified TAI removes the directionality aspect of the TAI metric by taking the absolute value of the segment mean.

All cross-sample CIN metrics comparisons using Spearman’s correlation (20) were conducted using the base ‘stats’ R package (21) (Version 4.0.4) ‘cor’ function between the non-tumor (“Blood Derived Normal,” “Solid Tissue Normal,” “Bone Marrow Normal,” and “Buccal Cell Normal”) and tumor (“Metastatic,” “Primary Blood Derived Cancer,” “Primary Tumor,” “Recurrent Tumor,” “Additional - New Primary,” and “Primary Blood Derived Cancer - Peripheral Blood”) samples. All heatmaps were generated using the ‘ComplexHeatmap’ R package (Version 2.9.3) (22) and clustered with the “complete’ method by “Euclidean distance.”

For analyses that compare CIN metrics by clinical characteristics, we used the TNM (tumor stage, node, and metastasis) staging provided by TCGA in the “ajcc_pathologic_m” (except for ACC where we used “ajcc_clinical_m”), “ajcc_pathologic_n,” and “ajcc_pathologic_t” attributes using the ‘GDCquery_clinic’ function. These comparisons were made across the 22 cancer types with corresponding data (i.e., all but Glioblastoma (GBM), Acute myeloid leukemia (LAML), Ovarian cancer (OV), Thymoma (THYM), Uterine carcinoma (UCS), Diffuse large B-cell lymphoma (DLBC), Pheochromocytoma and paraganglioma (PCPG), Uterine corpus endometrial carcinoma (UCEC), Sarcoma (SARC), Prostate adenocarcinoma (PRAD), and Brain lower grade glioma (LGG)). In addition, we conducted the Mann Whitney Wilcoxon Test (23,24) for comparisons between CIN metrics and the staging variables using the ‘rstatix’ R package (Version 0.7.0) (25) ‘wilcox_test’ function. We used Bonferroni-corrected p-values to account for multiple hypothesis testing, and we considered corrected p-values of less than 0.05 to indicate significant CIN metric scores for the TNM staging analyses.

For analyses comparing CIN metrics by sex for each cancer, we used the ‘gender’ (i.e., biological male or female) information provided by TCGA for each sample. We compared 27 cancer types because TCGA projects OV, PRAD, UCEC, UCS, Testicular germ cell tumors (TGCT), Cervical squamous cell carcinoma, and endocervical adenocarcinoma (CESC) occur predominantly in one biological sex. We conducted a Mann Whitney Wilcoxon test between CIN metrics and the sex variable using the ‘rstatix’ R package (Version 0.7.0) ‘wilcox_test’ function and multiple hypothesis tests corrected using the Bonferroni method. We plotted raincloud plots of these multiple hypothesis corrected p-values by CIN metric using the ggplot2 (Version 3.3.5), ggpubr (Version 0.4.0), PupillometryR (Version 0.0.4), and gghalves (Version 0.1.1) R packages for the top cancer identified by each CIN metric.

## 3. Results

We compiled masked CNV data for 11,124 patients across all TCGA projects (n=33 cancer types, 22,629 samples) and applied the six CIN inference calculations in the CINmetrics R package to each cancer data set (Figure 1).

**Figure 1.**
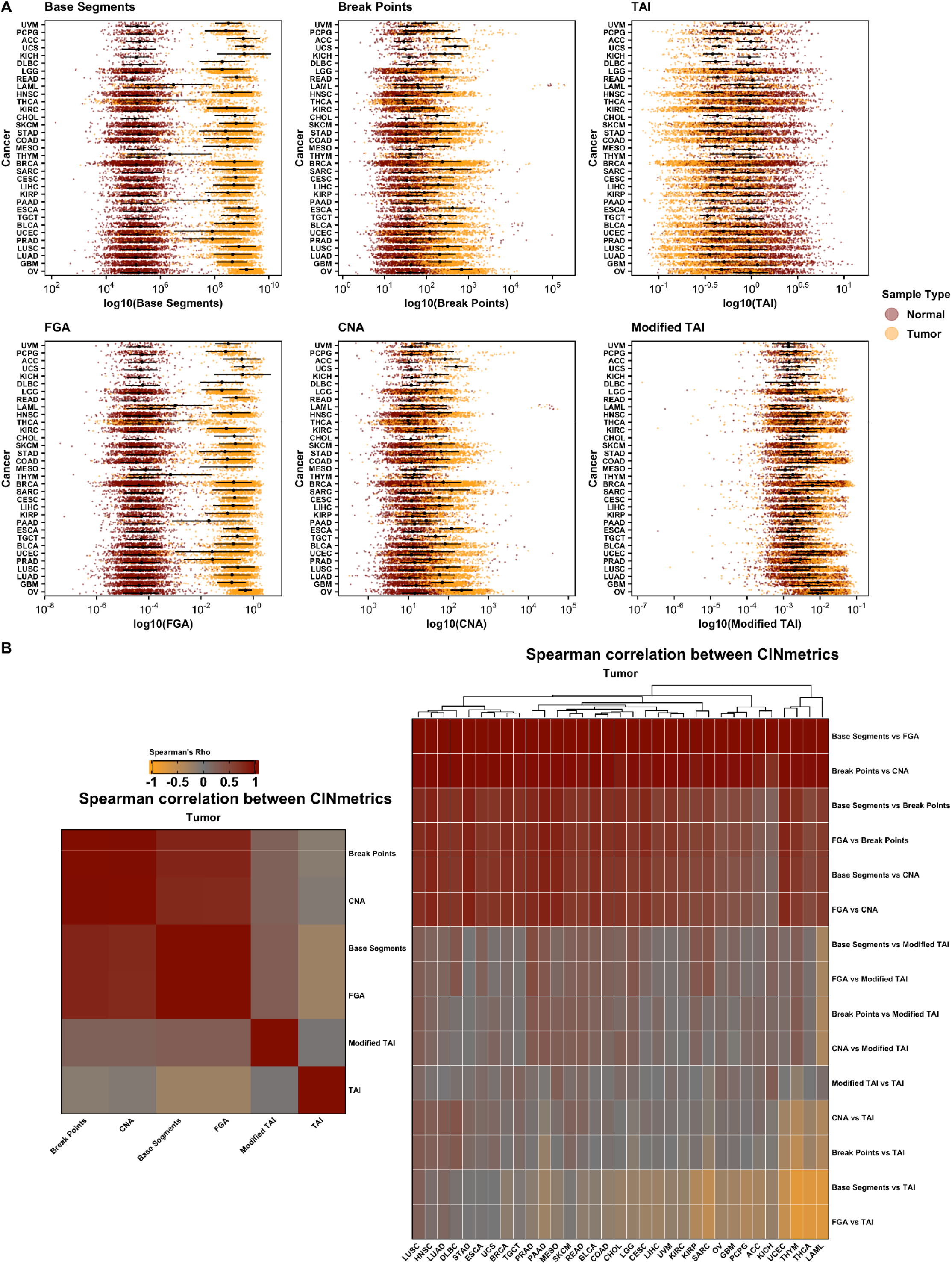
CIN metrics by cancer type for tumor and matched non-tumor samples (A) and Spearman’s correlation between tumor CIN metrics by cancer type (B). CIN, chromosomal instability.

Figure 1A shows the distribution of CIN scores across normal and tumor samples. Within each CIN “type” - structural (Base Segments, FGA), numerical (CNA, Break Points), or overall (TAI, Modified TAI) - the distribution pattern is consistent. However, there are noticeable differences across these types. For example, structural CIN metrics show a more pronounced distinction between normal and tumor samples compared to numerical CIN metrics. This is likely due to comparative genomic hybridization arrays used in TCGA to measure CNVs which are more biased towards detecting numerical CIN than structural CIN (26). Thus, one should be careful in evaluating genomic instability based on the choice of the metric. Subsequently, we conducted a Spearman’s correlation to analyze the relationship between each CIN metric across all tumor patient samples irrespective of the cancer type and within each cancer type. The results are depicted in Figure 1B, which further emphasizes the point that different “types” of CIN metrics capture varying aspects or patterns of genomic instability. Overall, Base Segments and FGA showed the most separation between tumor and non-tumor samples by cancer type, and as expected, the directionality of TAI for tumor compared to non-tumor samples is opposite of the other metrics (Figure 1A). Additionally, the tumor type impacts how well each CIN metric correlates with the others. For example, Base Segments and FGA show a positive correlation across all cancer types. However, when comparing Base Segments to TAI and FGA to TAI, the correlation varies based on cancer type. For both comparisons, there is a positive correlation in Lung squamous cell carcinoma (LUSC), Lung adenocarcinoma (LUAD). There is no correlation in Stomach adenocarcinoma (STAD), Esophageal carcinoma (ESCA), Uterine carcinoma (UCS), and a negative correlation in Thymoma (THYM), Thyroid carcinoma (THCA), Acute myeloid leukemia (LAML).

Next, we determined if each CIN metric is significantly associated with clinical characteristics by tumor type. For these analyses, we used the TNM (tumor stage, node, and metastasis) staging provided by TCGA for each of the 22 cancer types based on TNM data availability (Figure 2). We found that breast invasive carcinoma (BRCA; significant for Base Segments, FGA, Break Points, and CNA; Figure 2A-D), rectum adenocarcinoma (READ; significant for Break Points, and CNA; Figure 2C-D), head and neck squamous cell carcinoma (HNSC; significant for CNA; Figure 2D), and LUSC (significant for TAI; Figure 1E) each had at least one CIN metric significantly associated with tumor stage (Mann Whitney Wilcoxon Test <0.05 after Bonferroni correction). However, only colon adenocarcinoma (COAD; significant for Base Segments, FGA, Break Points, and CNA; Figure 2A-D) had CIN metrics significantly associated with metastases. COAD (significant for Base Segments and FGA; Figure 2A-B), kidney renal papillary cell carcinoma (KIRP; significant for Break Points and CNA; Figure 2C-D), and lung adenocarcinoma (LUAD; significant for Modified TAI; Figure 2F) all had at least one CIN metric associated with nodal involvement (Mann Whitney Wilcoxon Test <0.05 after Bonferroni correction).

**Figure 2.**
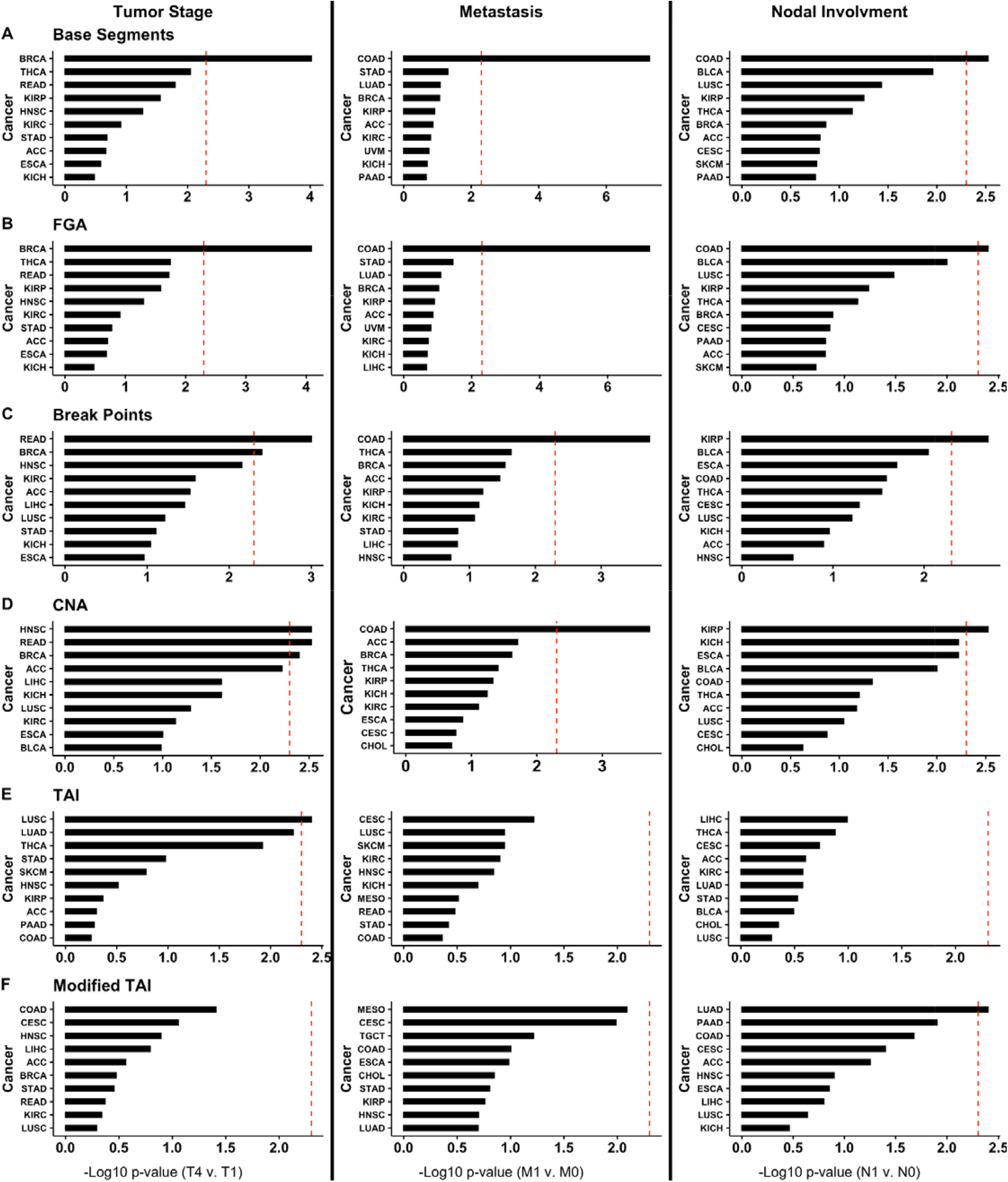
Top ten cancers with the lowest Bonferroni-corrected p-values for (A) Base Segments, (B) FGA, (C) Break Points, (D) CNA, (E) TAI, and (F) Modified TAI association with tumor stage (T4 compared to T1), metastasis (M1 compared to M0), and nodal involvement (N1 compared to N0). TAI, total aberration index; CNA, copy number abnormality; FGA, fraction genome altered.

Finally, we compared CIN metrics by sex for each cancer by using the ‘gender’ (i.e., biological male or female) information provided by TCGA for each patient for the 27 cancer types with cases in both sexes (Figure 3). We found that HNSC CIN was significantly different between the sexes based on the Base Segments, FGA, Break Points, CNA, and Modified TAI metrics. However, esophageal carcinoma (ESCA) was significantly different between the sexes based on the Break Points and CNA metrics (Figure 3A-B), THCA based on the CNA metric (Figure 3A), and GBM (Figure 3A-B), COAD (Figure 3A-B), kidney renal clear cell carcinoma (KIRC), READ, LUAD, and LUSC based on the Modified TAI metric (Mann Whitney Wilcoxon Test <0.05 after Bonferroni correction, Figure 3A).

**Figure 3.**
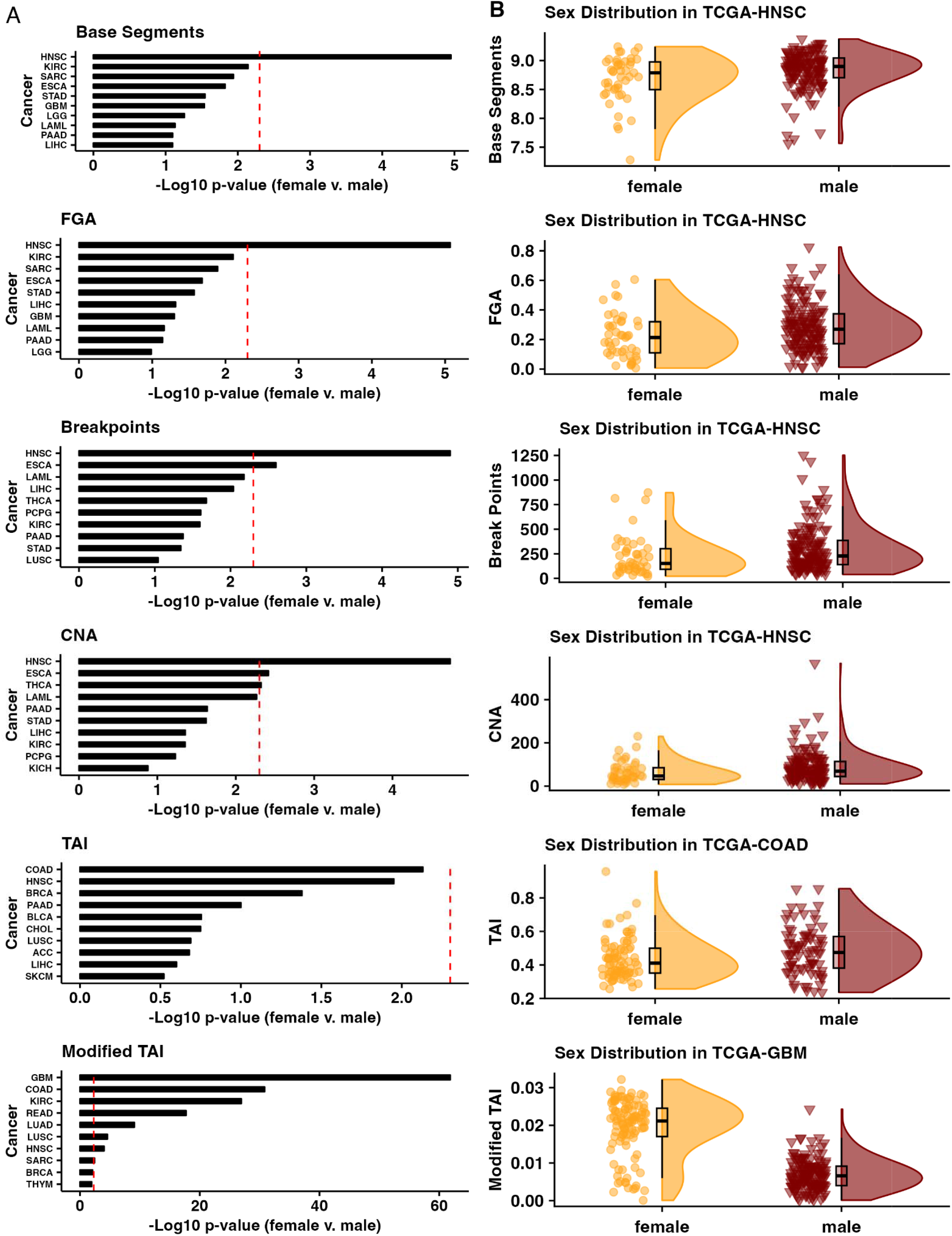
(A) Top ten cancers with the lowest Bonferroni-corrected p-values between the sexes for each CIN metric and the (B) distribution of CIN metric scores for the most different tumor cohort by sex for each CIN metric. CIN, chromosomal instability.

## 4. Discussion

Here we evaluated how the TAI, Modified TAI, CNA, Base Segments, Break Points, and FGA copy number aberration CIN surrogates (ie., names..) compared across TCGA cohorts by assessing each across tumor types, how they associate with tumor stage, metastasis, and nodal involvement, and with respect to patient sex. We found that the tumor type impacts how well any two given CIN metrics correlate. While we also identified overlap between CIN metrics regarding their association with clinical characteristics (e.g., CIN was significantly associated with tumor stage in BRCA for 4 of the 6 metrics) and patient sex (e.g., CIN was significantly different between the sexes in HNSC for 5 of the 6 metrics), there was not complete agreement between metrics. We identified several cases where only one CIN metric was significantly associated with a clinical characteristic or patient sex for a given tumor type (e.g., Modified TAI was the only CIN metric significantly associated with nodal involvement for LUAD). Therefore, caution should be used when describing CIN based on any one metric or when comparing across studies (27).

A recent study of 1,421 samples from 394 tumors across 22 tumor types demonstrates that somatic copy number aberrations in cancer are both pervasive (i.e., occurring at least once in 99% of tumors) and dynamic (i.e., more than 20% of the genome was subject to subclonal somatic copy number aberrations in 45% of tumors) (28). Generally, CIN has been associated with distinct cancer etiologies and progression, patient prognosis, drug efficacy, and drug resistance in tissue- and tumor-specific manners (4–11,29,30). For example, van Dijk et al. (30) found that variation in the chromosomal copy number within a tumor (CIN heterogeneity) was strongly associated with poor survival in patients with solid tumors such as breast cancer, lung cancer, and colorectal cancer. However, as we show, the choice of metric used to calculate CIN can affect the measurement of such variation. In (29), Lukowet al. discuss the role of aneuploidy in cancer drug resistance by either overexpression of genes that suppress DNA repair, promote cell growth, or through the accumulation of mutations in genes that encode DNA repair enzymes. Thus, in such cases measuring not only the CIN but the region where it occurs might be more useful for predicting drug resistance. While this indicates that CIN may be a promising biomarker of patient prognosis and drug response, a thorough understanding of how to measure and interpret CIN is critical. Our study further underscores the need to be specific about how and when during the disease course CIN is calculated and that patient characteristics like sex may impact or be associated with such metrics. For example, a previous study has reported that 73.1% of HNSC patients were male (31), and Park et al. show that males are at a 2.9-fold increased risk of HNSC, independently of tobacco and alcohol consumption (32). Additionally, multiple studies have shown an association between increased CIN and HNSC risk (33,34), all of which is in agreement with our report of higher CIN (for 5 out of 6 metrics) in males with HNSC.

The metrics used in this study reflect distinct aspects of CIN, such as numerical aberrations, structural aberrations, or whole genome instability, and each aspect may have different biological implications. For example, numerical aberrations might lead to a higher degree of genetic diversity within a tumor, providing a larger pool of genetic variants for natural selection to act upon. This could accelerate tumor evolution and adaptation, potentially leading to more aggressive or treatment-resistant cancers. Whereas, structural aberrations might disrupt specific genes or regulatory elements, leading to more targeted effects on cell function (35–37).

There are several limitations to this study. The first is that these CIN metrics are calculated based on one genomic profile generated from a tumor, or a tumor sample, at a given time. Tumors are often very heterogeneous, including across time, so this provides only a snapshot of a dynamic system (27). Recent studies have underscored that some tumor types have a strong correlation between CIN and metastasis that may be associated with the timing of copy number aberrations occurring during tumor development as well as the tissue of origin (38). Additionally, the data used to calculate the scores in this study are array-based intensity scores from bulk profiles (18). High-throughput sequencing and single-cell technologies will continue to allow for more comprehensive profiling and provide an opportunity for future studies to characterize cancer CIN more precisely (39). While we examined how clinical (e.g., tumor stage) or patient (e.g., sex) associated with metrics of CIN, limited sample numbers preclude examining many factors simultaneously across all cancers. Finally, this brief report does not investigate how different causes of CIN may influence these metrics.

## 5. Conclusion

In this study, we evaluated 6 different CIN metrics (TAI, Modified TAI, CNA, Break Points, Base Segments, and FGA) present in the literature, across 33 cancers present in TCGA. We find that the tumor type significantly impacts the correlation between any two given CIN metrics. While there was an overlap between CIN metrics associated with clinical characteristics and patient sex, there was not complete agreement between metrics. CIN is a complex and multifaceted phenomenon that cannot be fully captured by a single CIN metric; therefore, we caution against using a single metric as a proxy of CIN, particularly if developing CIN as a proxy for drug response or cancer progression for clinical settings.

## Acronyms

CIN: Chromosomal Instability
TCGA: The Cancer Genome Atlas
TAI: Total Aberration Index
CNA: Copy Number Abnormality
FGA: Fraction of Genome Altered
CNV: Copy Number Variation
TNM: Tumor stage, node, and metastasis
GBM: Glioblastoma
LAML: Acute myeloid leukemia
OV: Ovarian cancer
THYM: Thymoma
UCS: Uterine carcinoma
DLBC: Diffuse large B-cell lymphoma
PCPG: Pheochromocytoma and paraganglioma
UCEC: Uterine corpus endometrial carcinoma
SARC: Sarcoma
PRAD: Prostate adenocarcinoma
LGG: Brain lower grade glioma
TGCT: Testicular germ cell tumors
CESC: Cervical squamous cell carcinoma, and endocervical adenocarcinoma LUSC Lung squamous cell carcinoma
LUAD: Lung adenocarcinoma
STAD: Stomach adenocarcinoma
ESCA: Esophageal carcinoma
THCA: Thyroid carcinoma
BRCA: Breast invasive carcinoma
READ: Rectum adenocarcinoma
HNSC: Head and neck squamous carcinoma
COAD: Colon adenocarcinoma
KIRP: Kidney renal papillary cell carcinoma
KIRC: Kidney renal clear cell carcinoma

## Acknowledgments

We would like to thank the members of the Lasseigne Lab for their critical and constructive feedback.

## Notes

**Data Availability Statement** The data that support this study are openly available at the Genomic Data Commons (GDC) data portal (https://portal.gdc.cancer.gov/) as ‘TCGA Level 3’ data. In addition, all analyses associated with this paper are on GitHub (https://github.com/lasseignelab/CINmetrics_Cancer_Analysis) and available at https://zenodo.org/record/7942543#.ZGPu8OzMJ4A.

**Funding Statement** ST, VHO, JLF, and BNL were supported by R00HG009678. ST, VHO, TMS, JLF, and BNL were supported by UAB funds to the Lasseigne Lab.

**Conflicts of Interest** The authors have no conflicts to declare.

https://zenodo.org/record/7942543#.ZGPu8OzMJ4A

### Competing Interest Statement

The authors have declared no competing interest.

### Summary of Updates

We have added additional information placing our results into the context of the field in the Discussion. We also added an acronym list and additional labels on Figure 2 to improve readability.

